# Induction of Activity Synchronization among Primed Hippocampal Neurons out of Random Dynamics is Key for Trace Memory Formation and Retrieval

**DOI:** 10.1101/757229

**Authors:** Yuxin Zhou, Liyan Qiu, Haiying Wang, Xuanmao Chen

## Abstract

Memory is thought to be encoded by sparsely distributed neuronal ensembles in memory-related regions. However, it is unclear how memory-eligible neurons react incrementally during learning to encode trace fear memory, and how they respond to cues to retrieve the memory. We implemented fiber-optic confocal fluorescence endoscopy to directly visualize calcium dynamics of hippocampal CA1 neurons in freely behaving mice, which were subjected to trace fear conditioning. Here we report that the overall activity levels of CA1 principal neurons showed a right-skewed lognormal-like distribution. A small portion of highly active neurons (termed Primed Neurons) exhibited high sensitivity to sensory stimuli and marked activity plasticity. The Primed Neurons maintained random activity status for at least 5 hours in multiple contexts, including those prior to training and prior to recall. Repetitive training induced Primed Neurons to shift from random activity to a well-tuned synchronization. Importantly, the emergence of activity synchronization coincided with the appearance of mouse freezing behaviors. In recall, a partial synchronization among the same population of Primed Neurons was induced from originally random activity, which also coincided with mouse freezing behaviors. Additionally, training-induced synchronization facilitated robust calcium entry into individual Primed Neurons. In contrast, most CA1 neurons stayed silent and did not respond significantly to tone and foot-shock throughout the training and recall testing cycles. In conclusion, highly active Primed Neurons are preferably recruited to encode trace fear memory, and induction of activity synchronization among Primed Neurons out of random dynamics is critical for trace memory formation and memory retrieval.

**Significance Statement:**

- The overall activity levels of hippocampal principal CA1 neurons show a right-skewed lognormal-like distribution.
- A small portion of CA1 neurons (termed Primed Neurons) exhibit high activity and marked activity plasticity during learning and recall.
- Primed Neurons in the hippocampus are preferably recruited to encode trace fear memory, whereas the majority of CA1 neurons show little activity and little activity change during fear conditioning and recall testing.
- Induction of activity synchronization among Primed Neurons out of random dynamics is critical for trace memory formation and memory retrieval.
- Training-induced synchronization drastically increases calcium entry into Primed Neurons, which may promote memory consolidation.

## Introduction

It has been recognized that a portion of sparsely distributed neurons (coined engram cells^1,2^) in memory-related regions are recruited to encode memories^3,4^. However, it is unclear how engram cells react during learning to encode declarative memory, and how they respond to cues to retrieve specific memories. One prevailing model proposes that engram cells activate during learning to encode a memory and the same group of neurons re-activates during recall to retrieve the memory^2,5–7^. This model was developed mostly based on contextual fear conditioning experiments using nuclei staining^7^ or engram cell manipulation technology^2^. While the model helps establish the concept of engram cells and highlights the contribution of gene expression of engram cells to memory consolidation, it remains unclear how the engram cell population operates in synergy to construct the network basis of declarative memory. We hypothesized that the engagement of engram cells with memory does not directly arise from their silent state; rather, engram cells first become sensitized and stay in a “primed” state before participating in memory formation and before retrieving a memory. Emerging evidence has demonstrated that a subset of neurons in the lateral amygdala^8^ and the hippocampus^9^ with increased excitability “win the competition” to be selected to encode contextual fear memory^9–13^. These findings suggest that there is a hierarchy in recruiting neurons to encode memory, in accordance with the concept that sensitized or Primed Neurons are the preferable candidates to encode memory. However, most hippocampus-dependent memory studies were limited to investigating contextual fear memory, which associates spatial information of the surrounding environment with aversive stimuli, whereas the cellular or network mechanisms of trace fear memory formation have not been well elucidated.

Although the hippocampus is required for the formation of both spatial memory and semantic memory^28^, encoding mechanisms of two types of memory may be fundamentally distinct. We chose a trace fear conditioning behavioral paradigm, as this model provides manifold advantages to study declarative memory. Trace fear conditioning generates various memories, not only pertaining to spatial information of the environment, but also memories for the foot-shock event, such as the causal relationship and timing information of the tone and foot shock^14–16^. Acquisition of trace fear memory is considered a declarative task, which requires higher brain function including attention^17^ and the conscious awareness of the conditional stimulus (CS) - unconditional stimulus (US) contingency^18^. Multiple brain regions including the medial prefrontal cortex^18^, amygdala^18,19^, and the hippocampus participate in the acquisition of trace fear memory. Thus far, it is unknown how neurons in these regions behave before learning or react stepwise in response to CS-US paired training to form trace fear memory, nor how memory-holding neurons respond to a cue to retrieve a memory.

To directly visualize the real-time process of trace memory formation and retrieval, we implemented an *in vivo* calcium imaging technology, termed fiber-optic confocal (FOC) fluorescence endomicroscopy, in our investigation^20^. We combined the FOC calcium imaging system with trace fear conditioning in a time-locked manner. This combination approach enabled us to correlate mouse behaviors with *in vivo* calcium dynamics, which infers neuronal electrical activity or action potential back-propagation-induced calcium entry. Importantly, it also allowed for the detection of real time activity change of the same neuronal population in a deep-brain region, as these neurons engage in the process of memory formation in learning and respond to cues in subsequent recall. Here we show that the overall activity levels of CA1 principal neurons show a right-skewed lognormal-like distribution. High activity Primed Neurons are preferably recruited to encode trace fear memory, whereas the majority of CA1 neurons show little activity change during trace fear conditioning and recall testing. Further, induction of activity synchronization among Primed Neurons out of random dynamics is critical for trace memory formation and memory retrieval.

## Methods and Materials

### Mice

Calcium imaging and behavioral procedures were approved by the Institutional Animal Care and Use Committee at the University of New Hampshire and performed in accordance with their guidelines. C57Bl/6 mice and forebrain excitatory neuron-specific EMX1-Cre mice^21^ were bred and genotyped as previously reported ^22^. Data from 9 male and 6 female mice at ages 2-5 months were analyzed. We did not observe marked differences between males and females in developing activity synchronization and fear behaviors during training and recall. Presented data were pooled from both sexes. Mice were maintained on a 12-h light/dark cycle at 22°C and had access to food and water *ad libitum*. Mice were handled by investigators for 10 min and habituated to the hooking of a fiber optic-like wire to a pre-implanted cannula for 20 min for 3-4 days (one time per day) to allow them to adjust to investigators and adapt to the mounted fiber-optic micro-probe before starting the trace fear conditioning experiments.

### FOC imaging configuration

The FOC system contains a bundle of 8409 fiber optics (1 µm each) encased in a 300 nm micro-probe that conveys excitation light to the tissue and collects the emitted fluorescence from the tissue^20,23^. One end of the optic fibers encased in the micro-probe is inserted through an implanted cannula into a brain area to detect fluorescence signals, and the other hooks to the laser micro-objective, which connects to the imaging processing unit (Figure 1). Compared to direct mounting a miniature scope in the brain, inserting a fiber-optic microprobe into the brain can significantly reduce tissue damage and endure imaging stability. A pre-implanted cannula on the skull can hold the fiber-optic microprobe very tightly, which effectively minimizes the impact of animal locomotion on the imaging stability. The FOC imaging configuration doesn’t restrict free exploration and behaviors of mice.

**Figure 1.**
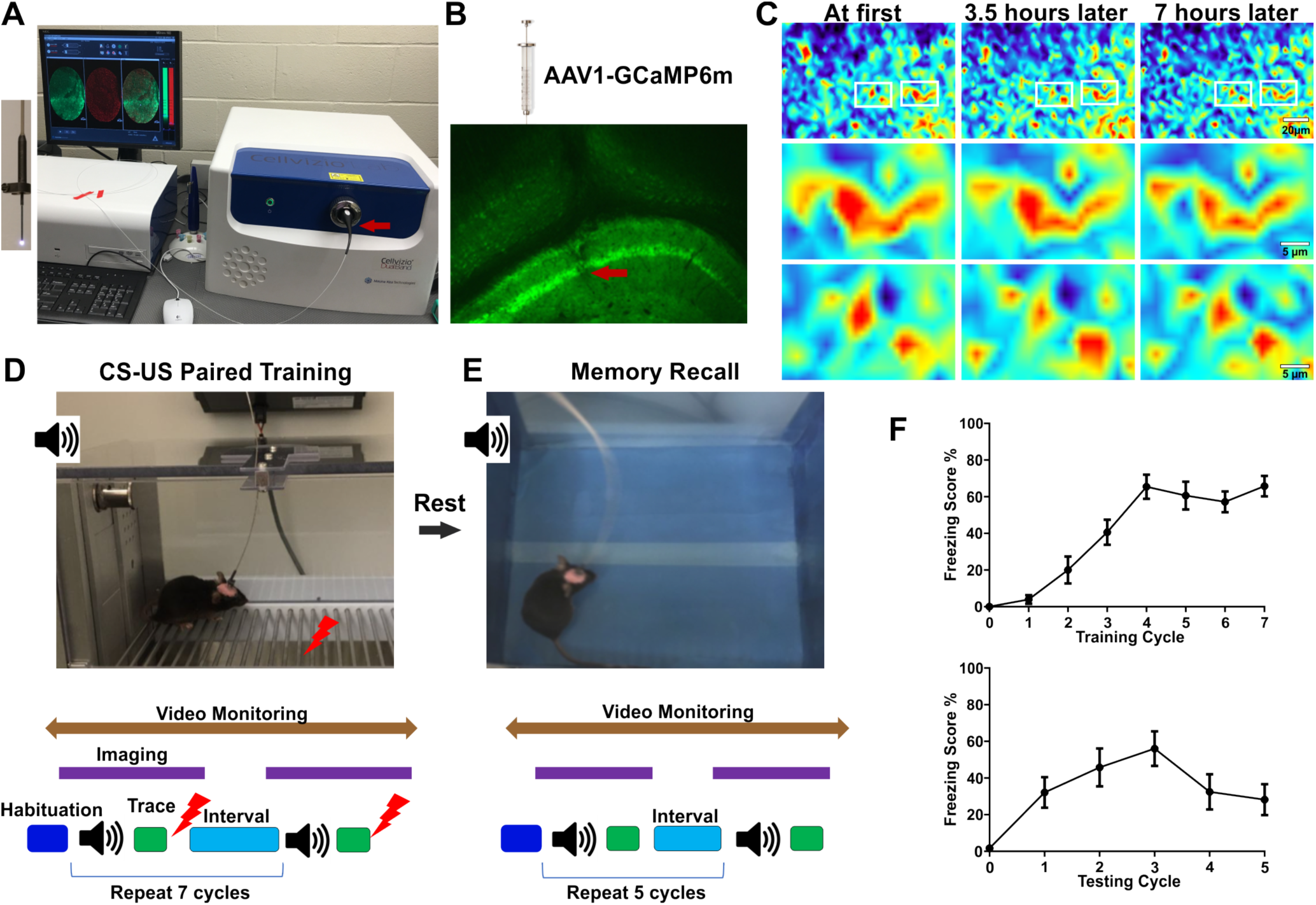
Configuration of *in vivo* FOC imaging system in freely behaving mice. (A) An imaging process unit of Cellvizio® Dual-Band 488/660 imaging system. A fiber-optic objective (red arrow) connects the imaging process unit with optic fibers. Left insert shows a fiber-optic microprobe. (B) AAV1-GCaMP6m or Cre-dependent AAV1-Flex-GCaMP6m were injected to the hippocampal CA1 region of C57Bl/6 mice or EMX1-Cre mice. Immunofluorescence staining shows GCaMP6m expression in the hippocampus with an arrow pointing the imaging location in the CA1 layer. Imaging locations could also be verified during imaging by a dense layer of neurons in the CA1. (C) Stability and single cell resolution of the FOC imaging. Representative images from the same animal demonstrate that the same neurons in the same location could be stably imaged for 7 hours. Single-cell resolution images were attained by the FOC imaging system (zoom in images in the bottom). (D) Top: a mouse carrying a FOC microprobe in a fear conditioning chamber. Bottom: learning paradigm of trace fear conditioning. (E) Top: recall testing environment, which was distinct from the learning context. Bottom: a paradigm of recall testing. Mice had 2-3 hours’ rest between training and the subsequent recall testing. (F) Freezing scores of mice in trace period during training (top) and during recall testing (bottom). n=9.

Because neuronal activity (synaptic activity and action potentials) leads to membrane depolarization and elevation of intracellular calcium in the cell body, GCaMP6m, a genetically-encoded calcium indicator, was expressed in neurons as a calcium sensor to monitor the activity-dependent calcium dynamics that reflect neuronal activity ^24^. GCaMP6m can be virally delivered and expressed at specific brain regions to label specific neuronal types when employing a Cre-loxP expression system together. We combined a deep-brain FOC imaging system (Cellvizio Dual-band 488/660 Neuropak^TM^) together with viral-mediated gene delivery to C57Bl/6 mice or to forebrain-specific EMX1-Cre mice to monitor the activity of excitatory neurons in CA1 regions of the hippocampus in freely-behaving mice.

### Mice surgery and virus injection

Mice were anesthetized with 1-3% isoflurane gas and mounted in a stereotaxic frame. The surgery was conducted as previously described^22,25^ with minor modifications. Briefly, a small skull window (coordination: AP: −1.85 to −1.95mm relative to the Bregma, ML: −1.4 mm relative to midline) above the dorsal CA1 region was made to allow installation of a special cannula (Mauna Kea Technologies, Paris, France) that accommodated an imaging FOC microprobe. The cannula was stably mounted on the top of the mouse skull using dental jet acrylic. Afterwards, we stereotaxically injected ∼0.5 µl AAV1 (purchased from Upenn Vector Core or Addgene) to mice through the cannula into the hippocampal CA1 region (coordination, AP: −1.9 mm to −2.0 mm; ML: −1.4mm; DV −1.4 mm). Our experiments were conducted first using C57Bl/6 mice injected with AAV1-GCaMP6m (purchased from Addgene, ID 100841). The hippocampal CA1 layer not only contains pyramidal neurons, but also harbors some interneurons, which was estimated to represent ∼5% total neurons in the mouse CA1^26–28^. To selectively visualize neuronal activity in excitatory principal neurons in the hippocampal CA1 region, we also injected AAV1-FLEX-GCaMP6m (purchased from Addgene, ID 100838) into the CA1 region of EMX1-Cre mice, whose Cre recombinase is expressed under the control of the promoter of forebrain-excitatory neuron specific EMX1 gene^21,22^. The approach led to selective detection of GCaMP6 imaging in principal neurons in the CA1 layer of the hippocampus. Because the expression of GCaMP6m in the two AAV vectors was both driven by the human Synapsin 1 promoter, very similar imaging pattern and behavioral results were obtained using C57Bl/6 mice (injected with AAV1-GCaMP6m) and EMX1-Cre mice (injected with AAV1-FLEX-GCaMP6m). We analyzed imaging data from 9 C57Bl/6 mice and 6 EMX1-Cre mice. In addition, this study focused on addressing the roles of Primed Neurons, and we combined several criteria to sort out Primed Neurons and Silent Neurons for analysis. These methods could eliminate the interference of interneurons, if detected occasionally in the imaging using C57Bl/6 mice, in our data interpretation. Therefore, we pooled the two datasets together to increase the statistical power of data analysis.

### Trace fear conditioning paradigm

3-4 weeks after surgery and AAV1 injection, mice were first positioned on a stereotaxic apparatus under isoflurane anesthesia. A CerboflexJ^tm^ Neuropak deep brain fiber optic microprobe (Mauna Kea Technologies, Paris, France) was hooked to a vertical micropipette guide of the stereotaxic apparatus that allowed positioning of the imaging probe and movement through the cannula until a bright neuronal image was seen. The imaging probe was tightly fixed to the cannula. Mice were then removed from the stereotaxic platform and put in a box with bedding, food, and water *ad libitum* for 1 hour to recover from isoflurane anesthesia. After completely waking up and regaining motor coordination, mice were then placed in a foot shock box (Med Associates Inc, Vermont) for experiments. The foot shock box was electrically connected to the FOC imaging system, which enabled the imaging procedure to be time-locked with the behavioral paradigm. A classic behavioral paradigm of trace fear conditioning was used. After 5-10 minutes of exploration and acclimation, a neutral tone, which is CS (3 KHz, 80 dB, 15 seconds) was delivered followed by a mild foot-shock, which was US (1 second, 0.7 mA) 30 seconds later. The CS-US pairing repeated 7 cycles (Figure 1), in which animals learned to associate CS with US and formed trace fear memory. After training, there was one additional imaging cycle without CS and US stimuli to monitor the basal neuronal activity in the absence of any stimuli. After completion of the learning procedure, mice were then put into a box with bedding, food and water *ad libitum* to rest for 2-3 hours before recall experiments. To avoid contextual cue to elicit contextual memory recall, we put mice into a novel environment distinct from the learning context (Figure 1), to monitor neuronal activity in response to tone. Mice were subjected to 5 cycles of tone stimuli (3 KHz, 80 dB, 5 seconds). Neuronal activity was monitored by the imaging probe. The trace fear conditioning and recall experiments were videotaped using a high-resolution monochrome camera and freezing behaviors were analyzed offline. Freezing was defined as complete lack of movement, except for breathing. Percent time spent freezing was derived by dividing the sum of freezing scores by the total time and multiplying the result by 100.

### Immunofluorescence staining

Mice were euthanized, and brain tissues were fixed with 4% paraformaldehyde and then subjected to fluorescence immunostaining using primary antibodies against GFP (1:500, Cat# A-11120, Thermo Fisher) followed by secondary antibody (conjugated with Alexa Fluor 488) staining to verify GCaMP6 expression and imaging location in the hippocampal CA1 region. Images were acquired with confocal microscope.

### Imaging data processing

We quantitatively analyzed neuronal activity throughout the experiments including during the trace memory learning period and in subsequent recall. Imaging data were acquired at 11.7 Hz by Cellvizio® Dual Band 488/660 imaging system (Mauna Kea Technologies, Paris, France) and analyzed off-line with IC Viewer 3.8 (Mauna Kea Technologies, Paris, France). Regions of interest (ROI) of calcium fluorescence for individual cells were manually circled with a diameter of 3-6 micrometers. Unstable imaging signals or motion artifacts were excluded through visual inspection of the ROI and fluorescence intensity. There were obvious distinctions between motion artifacts and real active neuronal signals. For example, motion artifacts randomly appeared in many neurons and they generally did not persist very long, while actual neuronal signals occurred in a small portion of neurons and their high activity lasted several hours. Unstable imaging or unreliable ROI signals were excluded for data analysis. The total number of cells recorded from each animal ranged from 50 to 200 neurons depending on AAV1 infection efficiency. Relative change in fluorescence intensity (%ΔF/F) was calculated by subtracting the normalized fluorescence intensity during baseline imaging from each data point during the imaging session. Throughout the manuscript, all imaging traces are raw traces of %ΔF/F, without filtering and digital processing. Thus, the imaging traces appear noisy, but an effective method to reduce noise is not yet available for this imaging system. Neuronal activity variance was calculated based on the relative change in fluorescence intensity (%ΔF/F). The intensity signals of ROIs were imported into Microsoft Excel, GraphPad Prism, and R programing language for further statistical analysis.

### Determination of tone-responsive neurons and foot shock-responsive neurons

We combined statistical analysis and visual inspection to determine tone-and foot shock-responsive neurons and calculate their percentage in the hippocampal CA1 region. We chose 3 second imaging trace (the relative change in fluorescence intensity - %ΔF/F) before and after the stimuli (tone or foot shock) and calculated their standard deviation (S-D). We defined the neurons as tone- or foot shock-responsive neurons, when the S-D of post-tone or post-foot shock %ΔF/F was more than two-fold higher than that of control period. Tone-responsive neurons and foot shock-responsive neurons were also verified by visual inspection of the imaging trace (%ΔF/F), if marked responding events appeared following a tone or a foot shock. Change-Point Analyzer software was also used to verify if a responsive event occurred following a tone or foot shock stimulus.

### Criteria to define Primed Neurons and Silent Neurons

We adopted multiple criteria to sort neurons and to define Primed Neurons and Silent Neurons: (1) The ratio of activity variance during the awake period relative to during the sleep period (Figure 3B). It was because virtually all neurons stayed inactive during sleep and had the lowest activity variance. The overall activity of CA1 neurons showed a right-skewed lognormal distribution with the ratios of high-activity neurons depicting the long-tail in the right (Figure 3B). We chose the top 10% right-skewed high-activity neurons as Primed Neurons. The selection of the top “10%” active was well-aligned with many previous reports^29,30^. We defined the bottom 70% with lowest activity variance ratio as Silent Neurons, and the intermediate 20% as Intermediate Neurons. The intermediate 20% may not be precise as a sharp line was yet to be identified to distinguish the Intermediate Neurons from Primed Neurons and Silent Neurons. (2) If their activity pattern changed after CS-US paired training (activity plasticity) - Primed Neurons exhibited strong activity plasticity, while Silent Neurons did not. (3) Sensitivity to tone and foot shock - Primed Neurons had the highest sensitivity, while Silent Neurons had little responses. Multiple cycles of CS-US paired training may slightly increase the activity of Intermediate Neurons. (4) If activity synchronization was formed after training – the Primed Neuron population developed activity synchronization after multiple cycles of CS-US paired training, while Silent Neurons did not. (5) Activity change in response to CS during memory recall – the activity pattern of Primed Neurons changed in responding to CS to develop partial synchronization, whereas that of Silent Neurons did not.

### Calculation of Synchrony Index (SI)

Let *y_it_* be the measured value from location *i* (*i* = 1, …, *I*) at time *t* for a neuron. Since the fluorescence signal detected by GCaMP6m slightly decreased with time in a linear manner due to a weak photobleaching effect, we subtracted the signal rundown by regressing the measure value on time. To be specific, we first fit a linear regression model 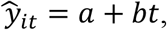 where

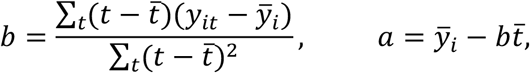

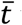 was the average time and 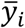 was the average measured value at location *i*. To remove the time effect, we used 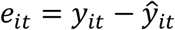 to calculate the Synchrony Index in the following way. For a pair of locations *i*, *j* = 1, …, *I*, calculate the absolute correlation coefficient

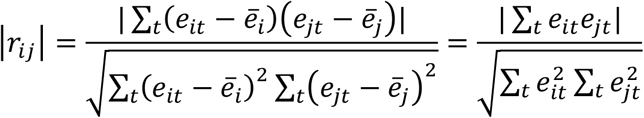

where the second equality was due to the fact that 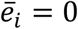 for all *i*.

We defined the Synchrony Index as the average of pairwise absolute correlation coefficient, i.e.,

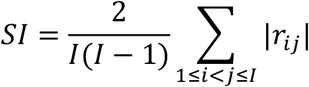

Here, we used the average of absolute correlation coefficients instead of the original correlation coefficients, because using the original correlation coefficients might otherwise fail to measure certain types of correlation.

### Calculation of Signal to Noise Ratio (SNR) for Variation

We also calculated SNR to characterize the fidelity of signal transmission and detection among neurons. Let *d_it_*= *y_i,t_* − *y_i,t-1_* and *d_jt_* = *y_j,t_* − *y_j,t-1_* for neuron *i* and *j*. Here *d_it_*’s can be interpreted as measurements on variations. A SNR was the ratio of the amount of variation caused by the variation from another neuron (signal) to the amount of variation caused by random noise. To define SNR, we first fit a linear regression model 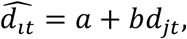 where

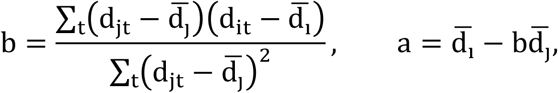

and 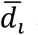 and 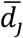 were the average values. The signal was measured by the sum of the squares of the regression (SSR)

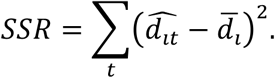

The noise was measured by the sum of the squares of the errors (SSE)

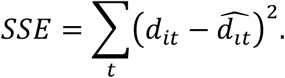

Thus, the SNR for a pair of neurons was defined as

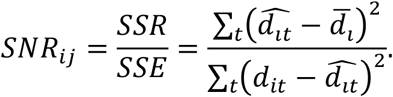

For a group of neurons, we defined the SNR to be

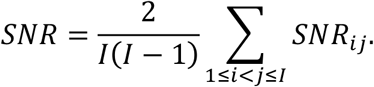

### Determination of “On-phase” and “Off-phase” of fluorescent calcium signals

We determined on-phase and off-phase of synchronized activity based on the activity dynamics of Primed Neurons during training, this is because Silent Neurons were constantly silent and did not display distinctions between on- and off-phases. The on-phase of Primed Neurons was chosen when synchronized activity patterns appeared in the trace period in the 5-7^th^ CS-US pairing cycle, whereas off-phase was defined as simultaneous silent phase (exhibiting low activity variance) in the trace period. The on-phase activity had at least 2-fold higher standard deviation than that of off-phase. In naïve condition - the Cy1, Primed Neurons did not have coherent activity in the trace period. The on-phase and off-phase were extrapolated by the same time frame of the on-phase and off-phase as determined in the training cycles, respectively. After multiple cycles of training, Primed Neurons had obvious synchronous events; the baseline values of the events were measured by the average normalized intensity prior to the events. In naïve conditions, when Primed Neurons demonstrated non-synchronous activity, the baseline value was determined by the average normalized intensity of the whole trace. The areas of on-phase and off-phase calcium signal density were calculated using ImageJ.

### Statistical analysis and graphic data presentation

Statistical analyses were conducted using paired or unpaired student t-test, or ANOVA test when appropriate. N.S. not significant, * *p*< .05, ** *p*< .01, \*\*\**p*< .001. A difference in data was considered statistically significant if *p* < .05, and values in the graph are expressed as means ± standard error of mean.

## Results

### *In vivo* FOC imaging configuration in combination with trace fear conditioning

FOC is a fiber-optic imaging system well-suited for accessing deep-brain regions to detect neuronal fluorescent signals in freely behaving mice^23,31,32^ (Figure 1A). To directly visualize neuronal activity in the hippocampal CA1 region during trace memory formation, we first virally delivered and expressed GCaMP6m, a genetically encoded calcium indicator^33^ carried by adeno-associated virus vector (AAV1) to the hippocampal CA1 region of mice (Figure 1B). The GCaMP6-imaging approach can simultaneously monitor calcium dynamics of a population of neurons (50-200 neurons) in freely behaving mice at the single cell resolution (Figure 1C). Imaging stability was critical for *in vivo* imaging in freely behaving animals, which enabled the visualization of the same population of neurons in different behavioral sections in real time. Due to the light-weight of the microprobe (Figure 1A) and improved implantation surgical skills, we were able to stably visualize the calcium signals of the same population of hippocampal neurons for 7 hours, despite vigorous exploration and jumping of animals during behavioral studies (Figure 1C).

We combined the FOC imaging approach with a trace fear conditioning paradigm (Figure 1 D&E) that trained mice to form an associative trace fear memory between an innocuous CS with an aversive US. A tone (CS) was followed by a mild foot-shock (US) 30 seconds later and the CS-US pairing repeated 7 cycles, in which animals stepwise learned to associate tone with foot shock, and developed a trace fear memory^14^. Typically, the freezing behaviors of mice emerged after 3 cycles of CS-US pairing (Figure 1F), indicating that animals have started to form an associative memory. Additional cycles of CS-US pairings reinforced the learning and increased the freezing score during the trace periods (Figure 1F). After conditioning, mice were allowed to rest or sleep for 2-3 hours. In recall, mice were put into a new environment and only tone was applied to test if an associative memory between CS and US had been formed (Figure 1E). During training, animal freezing behaviors became pronounced and freezing scores increased with cycles of CS-US paired training, while in recall animals’ freezing varied greatly from one trial to another (Figure 1F). These results were consistent with previous reports on trace fear conditioning^14–16^.

### Repetitive CS-US paired training drastically increases CS- and US-induced neuronal responses

The hippocampus is the information processing center of the brain, and sensory inputs are transmitted via naturally wired circuits into the hippocampus for information integration to form memories for knowledge, experience, events, and episodes^34^. We monitored calcium activity of hippocampal neurons on freely behaving mice. First, we evaluated how neurons in the hippocampal CA1 region naively responded to tone and foot shock without prior fear conditioning. Tone is an innocuous stimulus. Expectedly, animals did not respond strongly to tone on the first exposure, and tone-induced responses in these neurons did not sustain long either (Figure 2A). We did not identify many neurons naïvely responding to tone, and the percentage of tone-responsive neurons was estimated to be ∼0.6% (Figure 2D). Foot shock is an aversive stimulus; it was not surprising that the hippocampal CA1 region harbored more neurons that could naïvely and strongly respond to foot shock (Figure 2B). The percentage of foot shock-responsive neurons in the CA1 region was estimated to be ∼3.1% (Figure 2D). Hence, without fear conditioning, foot shock-induced neuronal responses outweighed tone-responses in the hippocampal CA1 region.

**Figure 2.**
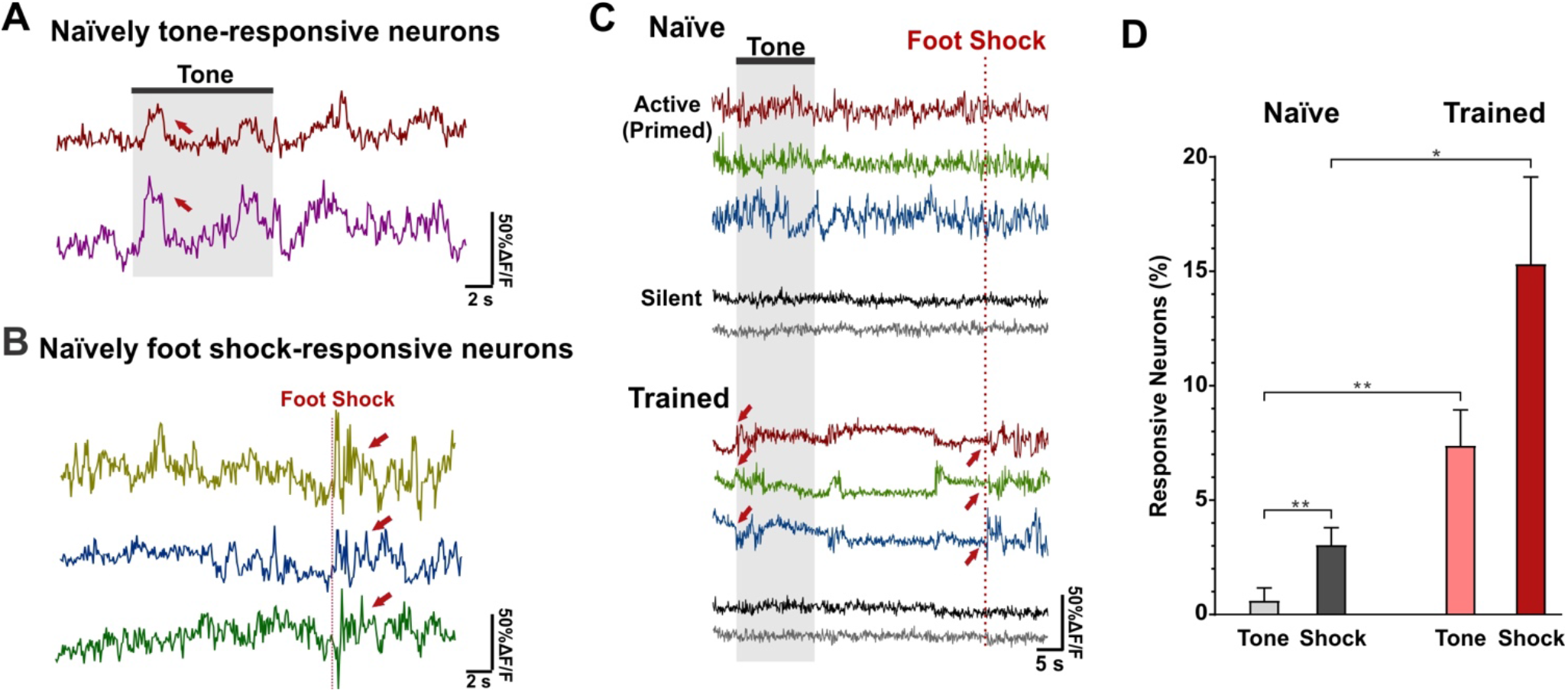
The percentage of tone- and foot shock-responsive neurons is drastically increased by CS-US paired training. (A) Representative fluorescence traces of two naïvely tone-responsive neurons. “Naïve” means the first exposure. Red arrows denote tone-evoked calcium signals. (B) Representative fluorescence traces of three naïvely foot-shock responsive neurons. Red arrows denote foot shock-evoked responses. (C) Representative neuronal responses under naïve condition and after being trained. Traces in same color represent the same neurons. Generally, most neurons including active neurons did not naïvely respond much to tone and foot shock on the first exposure. After multiple CS-US paired training, active neurons (color traces, also defined as Primed Neurons) were attuned to respond strongly to both tone and foot shock. However, Silent Neurons (bottom, two representative gray traces) with low activity variance did not change with training. (D) A bar graph shows the percentage of tone- and foot shock-responsive neurons, naïve versus after being trained. n = 5 animals, * p<0.05 with paired Student’s t-test.

We further observed that without fear conditioning, generally most neurons did not respond to tone or foot shock on the first exposure (Figure 2C). Mice were then subjected to multiple cycles of tone and foot shock (CS-US) paired trainings. Afterwards, tone-induced and foot shock-induced neuronal responses become obvious (Figure 2C). Interestingly, tone evoked much stronger neuronal responses, approaching a level close to the foot shock-elicited responses (Figure 2C), the percentage of tone-responsive neurons increased to ∼7.4%, and that of foot shock-responding neurons became ∼15.3% (Figure 2D). How did CS-US paired training cause these changes? We found that the increase came from randomly active neurons which did not markedly respond to tone or foot shock initially, but multiple cycles of CS-US paired training attuned them to strongly respond to both tone and foot shock (Figure 2C). In contrast, neurons that originally exhibited little activity variation (defined as Silent Neurons in the next section) still did not respond to tone or foot shock stimuli after training, and they did not contribute to the increase (Figure 2C). These data suggest that CS-US paired training recruited randomly active neurons, attuned them to respond to both tone and foot shock, and thereby drastically increased the percentage of CS-and US-induced neurons in the hippocampal CA1 region.

### The overall activity levels of principal CA1 neurons show a right-skewed lognormal-like distribution, and a small portion of active neurons (defined as Primed Neurons) can maintain high activity for 5 hours

The FOC imaging could stably visualize individual neurons’ calcium dynamics for up to 7 hours (Figure 1C), which allowed for continuous recording the real-time activity of individual neurons before and during training, and during recall testing as well as during sleep. First, we found that the activity dynamics of CA1 principal neurons varied drastically. Active (Primed) Neurons could stay active for several hours, while Silent Neurons constantly stayed silent even when subjected to repetitive training (Figure 3A). However, the activities of all neurons were reduced to a very low level during sleep. Surprisingly, a sleep epoch did not disturb the activity hierarchy among CA1 neurons: Primed Neurons regained high activity, and Silent Neurons still stayed silent after sleep (Figure 3A). Because the activity levels of all neurons, regardless of Primed or Silent Neurons, had no significant differences during sleep (Figure 3A&D&E, Supplemental Video 3), we used the activity variance of the relative imaging intensity (%ΔF/F) of each neuron during sleep as the “bottom-level” reference, and normalized the activity variance of individual neurons during the awake period to that of reference. This normalization removed individual signal detection variation and allowed for quantitative comparisons of activity levels among individual neurons. This normalization computation also yielded a pool of variance ratios at different time points for all individual neurons. High variance ratio denoted a high activity level while awake, whereas a ratio close to 1 indicated no activity difference between awake and sleep periods. We plotted a histogram of distribution probability for the ratios of all detected neurons (Figure 3B). The highest probability distribution occurred around a ratio of 1 (Figure 3B), confirming that most CA1 neurons were indeed silent. Further, training did not affect the general shape of the histogram plot, although sensory stimulation during training enhanced neuronal responses and slightly shifted the distribution plot to the right by a bin of 0.1 (Figure 3B). The histogram density of activity variance ratios was not a normal Gaussian distribution (Figure 3B). Rather, it exhibited a right-skewed distribution with high-activity neurons’ ratio filling the long tail in the right. Indeed, the histogram probability could be well fitted with a lognormal probability density function, 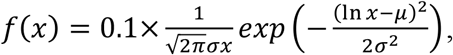 but not with a Gaussian function. Recently, right-skewed lognormal distributions have been described for several neurophysiology measurements including synaptic weight, network synchrony and firing rate of individual neurons^29,35^. Here, our results provide new evidence that the overall activity levels of CA1 principal neurons also exhibit a right-skewed lognormal distribution, supporting the concept that the pre-configured, right-skewed log-dynamics in the brain^29^ also include the overall neuronal activity in the hippocampus, and a highly active minority probably plays a dominant role in hippocampus-dependent memory formation.

**Figure 3.**
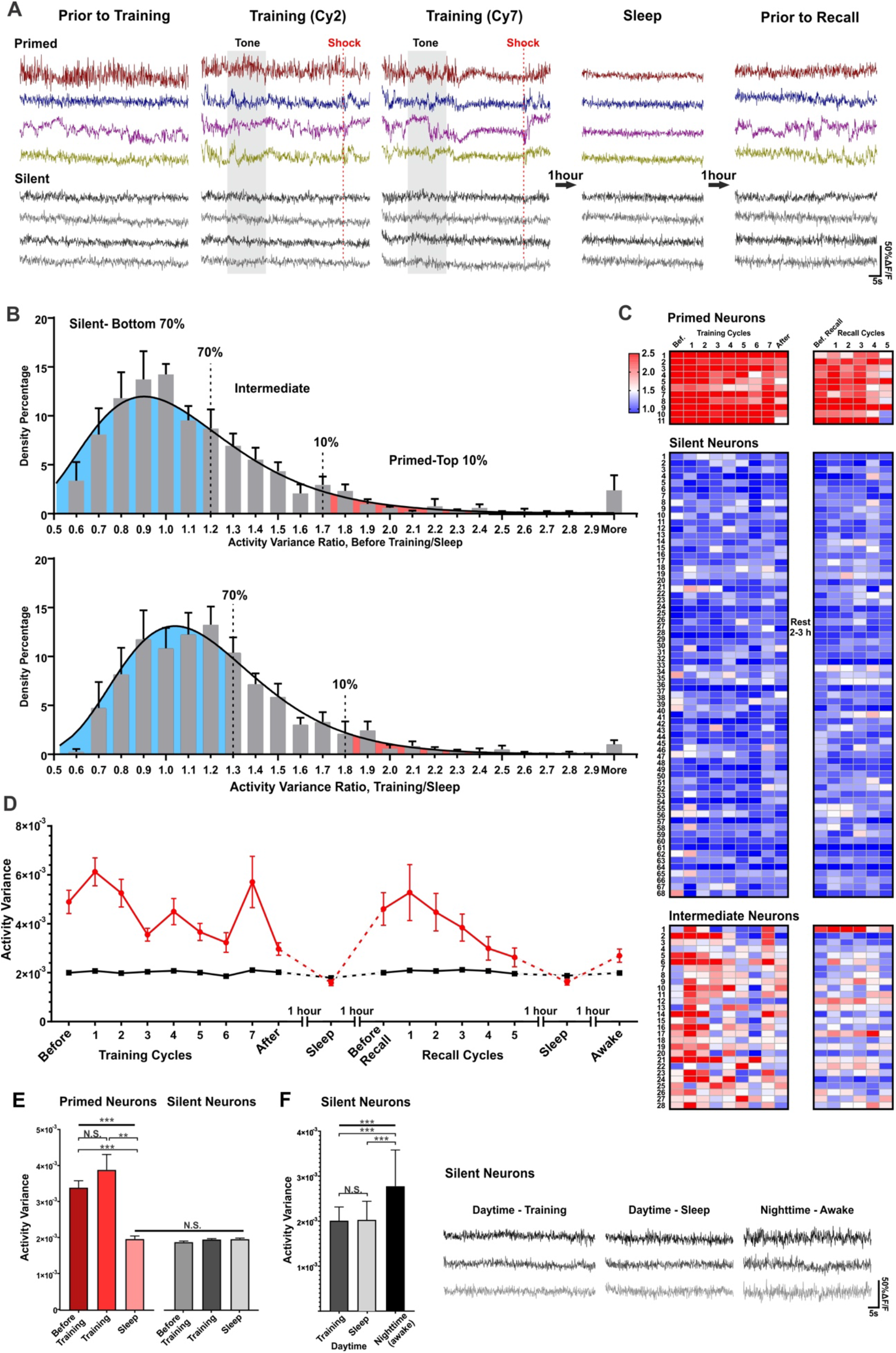
The overall activity levels of principal CA1 neurons show a right-skewed lognormal-like distribution. (A) Representative fluorescence traces of 4 Primed Neurons (color traces) and 4 Silent Neurons (grey traces) from one mouse at different time points. All traces in the figure have the same scale. (B) Histogram density of activity variance ratio of individual neurons. The variance ratio of individual neurons was calculated from its activity variance during awake (top, before training; bottom during training) divided by its activity variance during sleep. The histograms were well fit with a lognormal probability density distribution function, 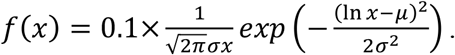 Top, before training, µ=0.02, σ=0.35, and bin width 0.1; Bottom, during training, µ=0.10, σ=0.29, and bin width 0.1. The data were computed from 800 neurons out of 6 mice. (C) Heat map of activity variance ratios during the course of training and recall testing. 11 Primed Neurons (top), 68 Silent Neurons (middle) and 28 Intermediate Neurons (bottom) from one animal are included in the heat map. Bef.: Before training. Activity variance ratios were activity variance at specific time points relative to that of sleep. Scale bar of the heat map is shown in the top left in which red colors indicate high-activity and blues mean low-activity. (D) Primed Neurons had much higher activity variance than Silent Neurons. Time-course of activity variance of representative 11 Primed Neurons and 11 Silent Neurons from one mouse. Primed Neurons had high activity variance before training, during training, and during recall testing, which could be maintained for 5 hours. During sleep, Primed Neurons and Silent Neurons had comparable low activity. (E) Activity variance of Primed Neurons (color columns) and Silent Neurons (grey columns) at different conditions. Primed Neurons had much higher activity variance during training as opposed to during sleep. One-way ANOVA, F (2, 75) = 13, ***, *p<*0.001. Post-hoc Tukey’s multiple comparisons test: Before Training Vs. Training, N.S., *p*=0.42; Before Training Vs. Sleep *p*<0.001, Training vs. Sleep *p*<0.01. Right 4 columns: Both Primed Neurons and Silent Neurons had low-level activity variances during sleep, and their activity variances had no differences with those of Silent Neurons before training and during training. One-way ANOVA, F (3, 463) = 1.589, *p*=0.1912, N.S. Post-hoc Turkey’s multiple comparison yielded no difference between any groups. Data were calculated from 89 Primed Neurons, and 527 Silent Neurons from 6 animals. (F) Silent neurons had similar activity level during training and during sleep in daytime, but their activity variance significantly increased at nighttime. One-way ANOVA with post-hoc Turkey comparison, F (2, 447) = 94, *p*<0.001. Post-hoc Turkey’s multiple comparison: Training Vs. Sleep, *p*=0.9536; Sleep Vs. Nighttime *p*<0.001; Training Vs. Nighttime, *p*<0.001. Representative traces of three Silent Neurons from one mouse at different times are shown in the right. Data were collected from 269 Silent Neurons out of 3 animals, whose images were acquired both daytime and nighttime.

Based on the variance ratio probability distribution (Figure 3B) and four other criteria including activity plasticity and sensitivity to sensory stimuli (described in Methods and Materials), we empirically classified principal CA1 neurons into three different groups. We defined neurons with the highest 10% variance ratios in the right as Primed Neurons, and the bottom 70% ratios in the left as Silent Neurons, and the remaining intermediate ∼20% as Intermediate Neurons (Figure 3B). Features of these three groups of neurons were summarized in Table 1. The first group of neurons was highly active prior to the training and sustained high activity levels during training (Figure 3A). They remained highly active during the subsequent recall testing 2-3 hours later (Figure 3C&D). The basal activity variance of Primed Neurons was the highest among the three groups (Figure 3C&D). Because this group of neurons were very sensitive to sensory stimuli and demonstrated activity plasticity during training and recall (see Figure 4&5), they were named Primed Neurons, meaning they were primed and prepared to participate in memory formation. Moreover, we also found that Primed Neurons could maintain highly active status (before training, during training, and after training) for 5 hours in a day (Figure 3C&D). It is worth mentioning that Primed Neurons might have already stayed active for a while before the imaging detection. The conclusion on activity duration of Primed Neurons were also in line with previous reports using a different imaging approach^30^.

**Figure 4.**
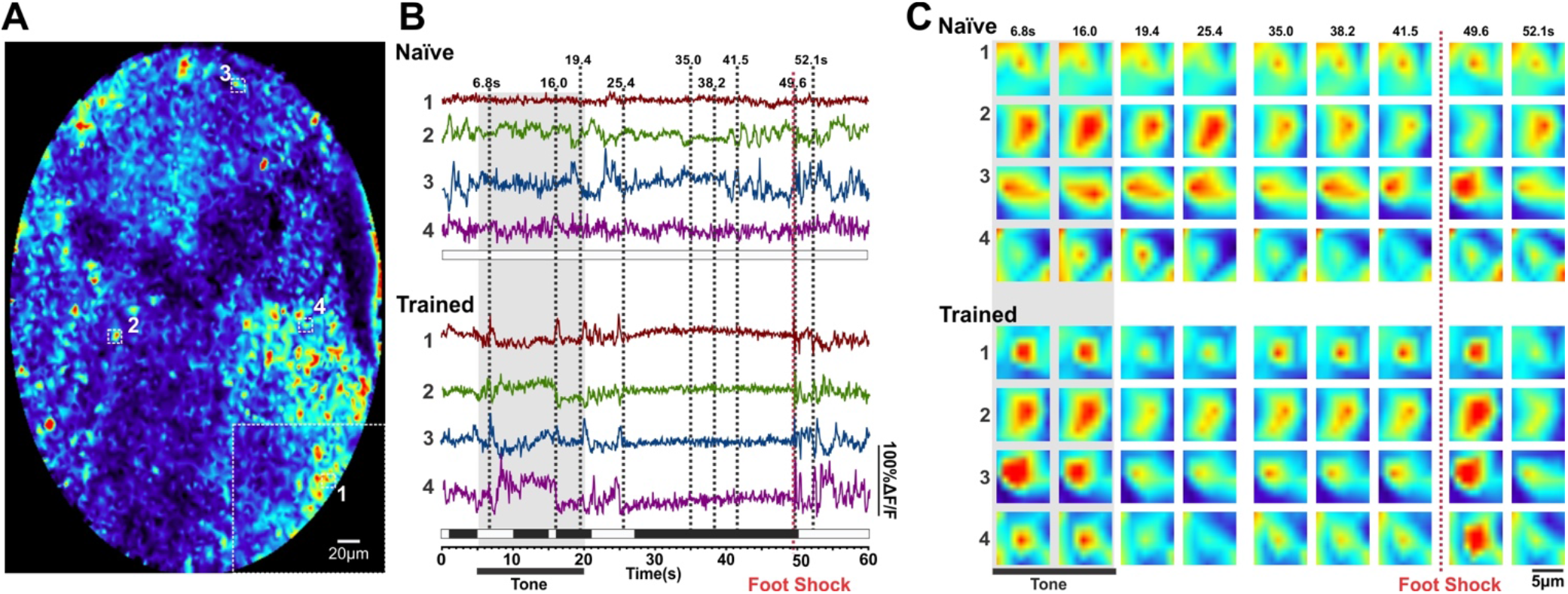
The activity dynamics of Primed Neurons are modified from random activity to synchronization by CS-US paired trainings. (A) Full endoscopic image with 4 representative Primed Neurons marked in their original locations. Neuron #1 had a different intensity scale from the other three neurons. (B) Representative fluorescence traces of 4 Primed Neurons. In naïve, the 4 Primed Neurons were randomly active and lacked activity coherence (top). However, their activity was modified by CS-US paired training to become more coherent (bottom). (C) Selected time-lapse images of 4 representative Primed Neurons in naïve condition (top) and after being trained (bottom). Dash black lines in (B) show different time points, when images were acquired. After being trained, the activity of the 4 Primed Neurons became synchronized: either simultaneously active or simultaneously inactive.

**Figure 5.**
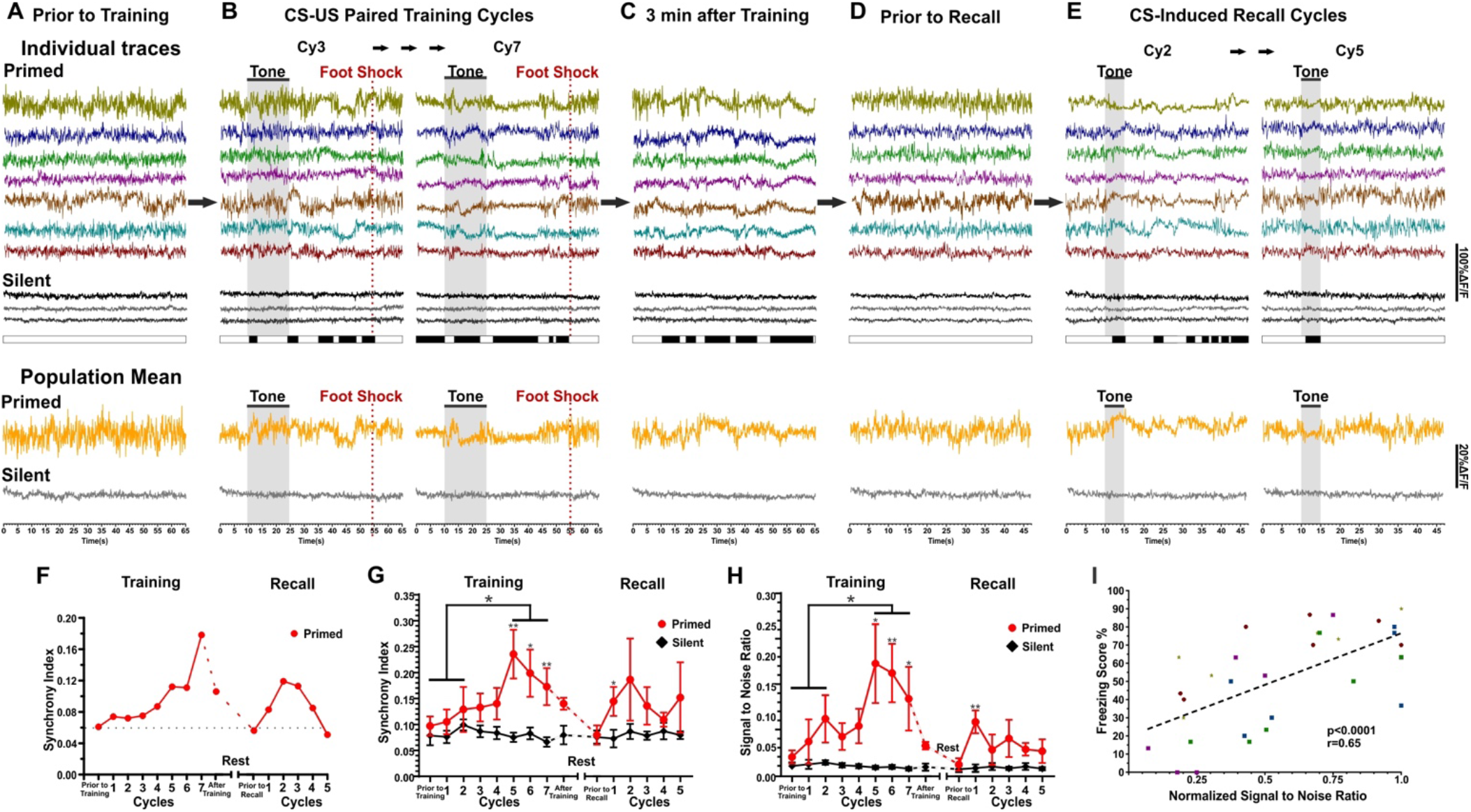
Induction of synchronization among Primed Neurons out of random activity is critical for trace memory formation and memory retrieval. (A) Representative fluorescent traces of 7 Primed Neurons (color traces) 3 min before training. Same color represents the same Neuron (for Panel A-E). Representative 3 Silent Neurons (gray traces) are shown below the Primed Neurons as a comparison. Beneath the traces are bars denoting animal’s behaviors, either freezing (black bars) or non-freezing (white bars). Population means of 7 Primed Neurons (orange traces) and 7 Silent Neurons (grey traces) are included in the bottom (Panel A-E). (B) The activity pattern of Primed Neurons was stepwise modified by repetitive training to form activity synchronization. The off-phase of synchronized activity strictly coincided with animal freezing behaviors. Cy3 and Cy7 denotes the 3^rd^ and the 7^th^ CS-US paired training cycles. Traces of the 7 Primed Neurons of all cycles are provided in Supplemental Figure 1. (C) The activity of Primed Neurons remained synchronized shortly after training. (D) After 2-3 hours rest, Primed Neurons returned to random activity prior to the recall. (E) Primed Neurons developed activity synchronization in response to tone during recall testing, which also coincided with freezing behaviors. (F) Synchrony Index of 7 Primed Neurons of one animal. Synchrony Index was defined as the average of pairwise absolute cross-correlation coefficient. (G) Primed Neurons had increased Synchrony Index with training. Summarized Synchrony Index of 38 Primed Neurons and 38 Silent Neurons out of 5 mice. During training, Primed Neurons in Cy5-7 had significantly higher Synchrony Indexes than these of not-trained cycles (prior to training-Cy0, Cy1 and Cy2). Comparison of not-trained cycles (Cy0, Cy1, and Cy2) with trained cycles (Cy5, Cy6 and Cy7) with unpaired Student’s t-test, *p*=0.008. Also, Primed Neurons in training Cy5-7 had significantly higher Synchrony Indexes than these of Silent Neurons: Cy5, *p*=0.01; Cy6, *p*=0.04; Cy7 *p*=0.02 with unpaired Student’s t-test. During recall, Cy1 had higher Synchrony Index than Silent Neurons, Cy1, *p*=0.03 with unpaired Student’s t-test. Silent Neurons’ Synchrony Indexes did not change significantly throughout training and recall cycles. Comparison of not-trained cycles (Cy0, Cy1, and Cy2) with trained cycles (Cy5, Cy6 and Cy7) with unpaired Student’s t-test, *p*=0.83. (H) Primed Neurons had increased SNR with training. SNR of Primed Neurons and Silent Neurons during training and recall testing. Comparison of not-trained cycles (Cy0, Cy1, and Cy2) with the trained cycles (Cy5, Cy6 and Cy7) with paired Student’s t-test, *p*=0.04. During training, Primed Neurons in Cy5-7 had significantly higher SNRs than Silent Neurons: Cy5, *p*=0.03; Cy6, *p*=0.008; Cy7, *p*=0.04 with unpaired Student’s t-test. In recall, Primed Neurons had higher SNR than Silent Neurons in Cy1, *p*=0.005 with Student’s t-test. The SNR of Silent Neurons did not change significantly throughout the training and recall cycles. Comparison of not-trained cycles (Cy0, Cy1 and Cy2) with trained cycles (Cy5, Cy6 and Cy7) with unpaired Student’s t-test, *p*=0.11. (J) The freezing scores of mice were correlated with normalized SNR of Primed Neurons during training. Data were collected from 5 animals. Different shape in the plot indicates data points from different animals. Linger regression of correlation analysis yielded r = 0.65 and *p*<0.0001.

**Table 1:**
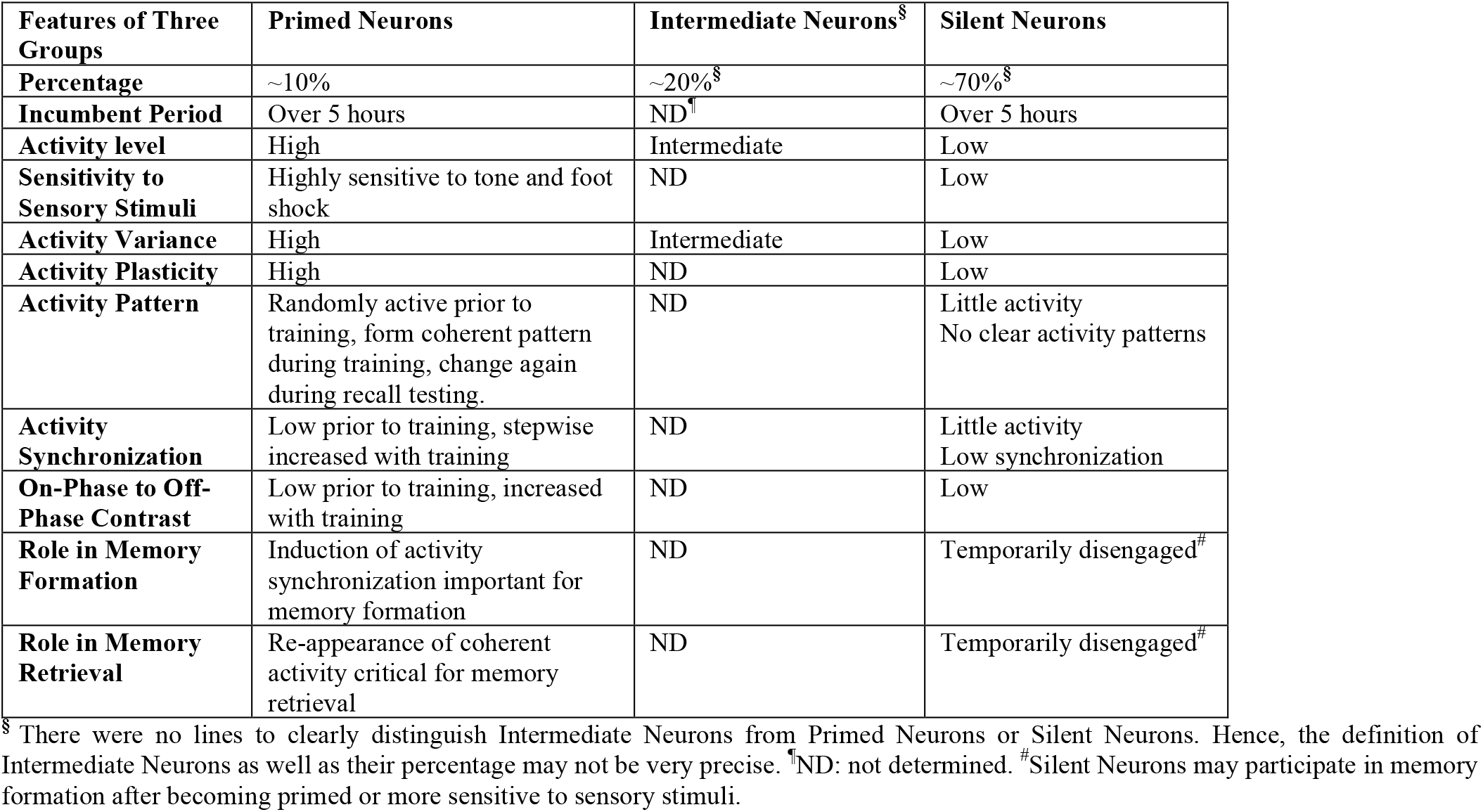
Features of Primed Neurons and Silent Neurons

| Features of Three Groups | Primed Neurons | Intermediate Neurons<sup>§</sup> | Silent Neurons |
| --- | --- | --- | --- |
| **Percentage** | ~10% | ~20%<sup>§</sup> | ~70%<sup>§</sup> |
| **Incumbent Period** | Over 5 hours | ND<sup>†</sup> | Over 5 hours |
| **Activity level** | High | Intermediate | Low |
| **Sensitivity to Sensory Stimuli** | Highly sensitive to tone and foot shock | ND | Low |
| **Activity Variance** | High | Intermediate | Low |
| **Activity Plasticity** | High | ND | Low |
| **Activity Pattern** | Randomly active prior to training, form coherent pattern during training, change again during recall testing. | ND | Little activity<br>No clear activity patterns |
| **Activity Synchronization** | Low prior to training, stepwise increased with training | ND | Little activity<br>Low synchronization |
| **On-Phase to Off-Phase Contrast** | Low prior to training, increased with training | ND | Low |
| **Role in Memory Formation** | Induction of activity synchronization important for memory formation | ND | Temporarily disengaged<sup>#</sup> |
| **Role in Memory Retrieval** | Re-appearance of coherent activity critical for memory retrieval | ND | Temporarily disengaged<sup>#</sup> |

<sup>§</sup> There were no lines to clearly distinguish Intermediate Neurons from Primed Neurons or Silent Neurons. Hence, the definition of Intermediate Neurons as well as their percentage may not be very precise. <sup>†</sup>ND: not determined. <sup>#</sup>Silent Neurons may participate in memory formation after becoming primed or more sensitive to sensory stimuli.

The intermediate group was moderately active. Their variance ratios were intermediate, lower than that of Primed Neurons but higher than that of Silent Neurons (Figure 3B). Intermediate Neurons became more active during training, however, their high-activity did not sustain long (Figure 3C - Bottom). It was somewhat ambiguous to distinguish this group from Primed Neurons and Silent Neurons. The third group was constantly silent and thereby named Silent Neurons. The majority of CA1 neurons were Silent Neurons (Figure 3B&C). Compared to Primed Neurons, the fluorescent traces of Silent Neurons were very flat and had the lowest basal activity variance (Figure 3A). Silent Neurons were unresponsive to tone and even to foot shock (Figure 3C, Figure 2C). Moreover, we did not detect marked activity modification among Silent Neurons by multiple cycles of CS-US paired training (Figure 3A-E). Silent Neurons stayed silent for more than 5 hours, even when stimulated with multiple tone or foot shock during training and during recall testing (Figure 3C&D). These results suggest that Silent Neurons may be temporarily disengaged from the engram network and contributed little to memory formation activity. Of note, the term “Silent Neurons” does not necessarily mean that these neurons were completely silent or lacking any action potential. They might have some background or noise-level electric activity, whose calcium spikes were probably too weak to be detected in the imaging system. Selection of region of interest (ROI) (3-6 µm in diameter) from endoscope images for data analysis were the same for all neurons, excluding the possibility of signal detection bias. To further rule out the possibility that the ROIs of Silent Neurons were not collected from cells, we compared the activity levels of Silent Neurons during the day with their nighttime activity levels. The activity variance of Silent Neurons significantly increased during nighttime (Figure 3F). This is because mice are nocturnal animals and become much more active during nighttime. The result suggests that Silent Neurons, while not active or easily excitable, do receive certain synaptic stimulation from other neurons, more so during nighttime than in daytime (Figure 3F).

### Induction of synchronization among Primed Neurons out of random dynamics by repetitive CS-US paired training is key for trace memory formation and memory retrieval

Next, we assessed the role of Primed Neurons in memory formation since they were highly sensitive to sensory stimuli. We first observed that in naïve status (the first CS-US paired exposure), Primed Neurons were randomly active and there was a lack of coherent activity among Primed Neurons (Figure 4, Supplemental Video 1). Nevertheless, after the animal was trained with multiple cycles of CS-US pairing, the activity dynamics of Primed Neurons became more coherent, either being active or silent simultaneously (Figure 4, Supplemental Video 2). We stepwise monitored and analyzed the activity dynamics of Primed Neurons during training and recall testing. Prior to the training, Primed Neurons were randomly active, while Silent Neurons generally stayed silent (Figure 5A). The activity pattern of Primed Neurons was modified by CS-US paired training in a stepwise manner to become coherent. Often, the activity modification of Primed Neurons emerged at the 3^rd^ training cycle and became more pronounced at the 5^th^-7^th^ cycle, when strong responses of Primed Neurons during tone and trace period were clearly visible (Figure 5B, Supplemental Figure 1). We calculated the synchrony index of Primed Neurons and found it to be low prior to the training, but it increased with cycles of CS-US paired training (Figure 5F&G). Importantly, the modification of Primed Neurons’ dynamics from irregularity to synchronization coincided with the appearance of mouse freezing behaviors (Figure 5B), with the off-phase (the inactive period) of the synchronization matching with its freezing behaviors. The activity synchronization of Primed Neurons and animal freezing persisted several minutes after training (Figure 5C), suggesting that the synchronization of Primed Neurons could recur shortly after training in the absence of CS and US. Signal to noise ratio (SNR) has been used in neuroscience to measure the reliability of neural information transmission^36^. SNR is the ratio of the amount of variation caused by the variation from another neuron (signal) to the amount of variation caused by random noise. We also calculated SNR to characterize the fidelity of signal transmission and detection (or synchronization) among Primed Neurons and among Silent Neurons. Figure 5H shows that the SNR of Primed Neurons increased with CS-US paired training. After training, the SNR gradually reduced. The SNR of Primed Neurons was correlated with animal freezing scores during training (Figure 5I). Additionally, population mean was used to describe burst synchronization of neuronal membrane potentials *in vitro*^37^, as synchronized activity does not subtract each other during averaging. We calculated the population mean of both Primed Neurons and Silent Neurons. Prior to training, there was no clear pattern among the Primed Neuron population (Figure 5A&B bottom). After CS-US paired training (Figure 5B, the 7^th^ cycle), a clear pattern of the population mean responding to tone and foot shock emerged, which coincided with the mouse freezing behaviors. Together, these data suggest that induction of activity synchronization among Primed Neurons is critical for trace memory formation.

In contrast, Silent Neurons did not exhibit activity modifications during trace fear conditioning (Figure 5B, Supplemental Video 1&2), and their Synchrony Index and SNR remained low and did not increase with repetitive CS-US paired trainings (Figure 5G-H). Moreover, their population mean did not change with training or display any coincidence with mouse freezing behaviors (Figure 5B). Together, these data demonstrate that Primed Neurons played a leading role in mediating trace fear memory formation, whereas Silent Neurons probably had little contribution to trace memory formation.

If formation of activity synchronization among Primed Neurons is critical for trace memory formation, then a synchronization would re-appear during recall testing. To test if animal freezing coincides with synchronization again in recall, we put animals into a novel environment distinct from the learning context and only introduced tone as a cue for recall testing. Interestingly, we found that the same population of Primed Neurons maintained highly active status, but their activity shifted back to random dynamics after the training (Figure 5D). They were randomly active prior to recall testing, but underwent marked activity modifications from random activity to synchronization in response to tone (Figure 5E). The synchrony index and SNR among Primed Neurons increased in recall compared to that prior to recall (Figure 5F&H). The emergence of activity synchronization among Primed Neurons also coincided with the appearance of mouse freezing behaviors (Figure 5E). The population mean of Primed Neurons exhibited marked change upon tone (Figure 5E-bottom). Nevertheless, Silent Neurons remained silent and displayed little activity modification upon tone (Figure 5E). The Synchrony Index and SNR among Silent Neurons (Figure 5G-H) and their population mean (Figure 5E-bottom) did not vary a lot during recall.

Note that the activity synchronization among Primed Neurons during recall was not as high as that during training (Figure 5G&H), therefore it was considered partial synchronization. Also, although during training, repetitive trainings led to higher synchronization and stronger freezing responses, memory recall induced by CS was somewhat stochastic and not every trial could successfully evoke freezing responses (Figure 5E). If tone failed to induce activity synchronization among Primed Neurons, animals did not display marked fear responses, even though their Primed Neurons stayed active (Supplemental Figure 1), suggesting that memory retrieval depends on the formation of activity synchronization of Primed Neurons, not simply neuronal activation. Together, these data support the conclusion that Primed Neurons play a leading role in encoding trace fear memory, and induction of activity synchronization out of originally random activity is critical for trace memory formation and retrieval.

### Training-induced synchronization condenses population activity into a narrow bursting frame, facilitating robust calcium entry into Primed Neurons

Principal neurons in the hippocampal CA1 region not only receive Schaffer-collateral synaptic projects from the CA3 region^34^, but also connect with each other with recurrent synapses^38^ as well as other projections. We reasoned that the calcium dynamics of Primed Neurons reflect the synchronization levels of the engram network and coherent synaptic inputs from other neurons. When a Primed Neuron received numerous asynchronous synaptic stimulations from the network to become electrically active, its calcium dynamics displayed a chaotic pattern (Figure 6A-Top). Conversely, when it received synchronized synaptic inputs from the network in a “bursting” pattern (Figure 6A-Bottom), its calcium dynamics would exhibit two phases: active “on”-phase and silent “off”-phase. The on-phase received synchronized stimuli from the network, leading to bursting activity of itself, whereas the off-phase got little stimulation and stayed silent (Figure 6A-Bottom).

**Figure 6.**
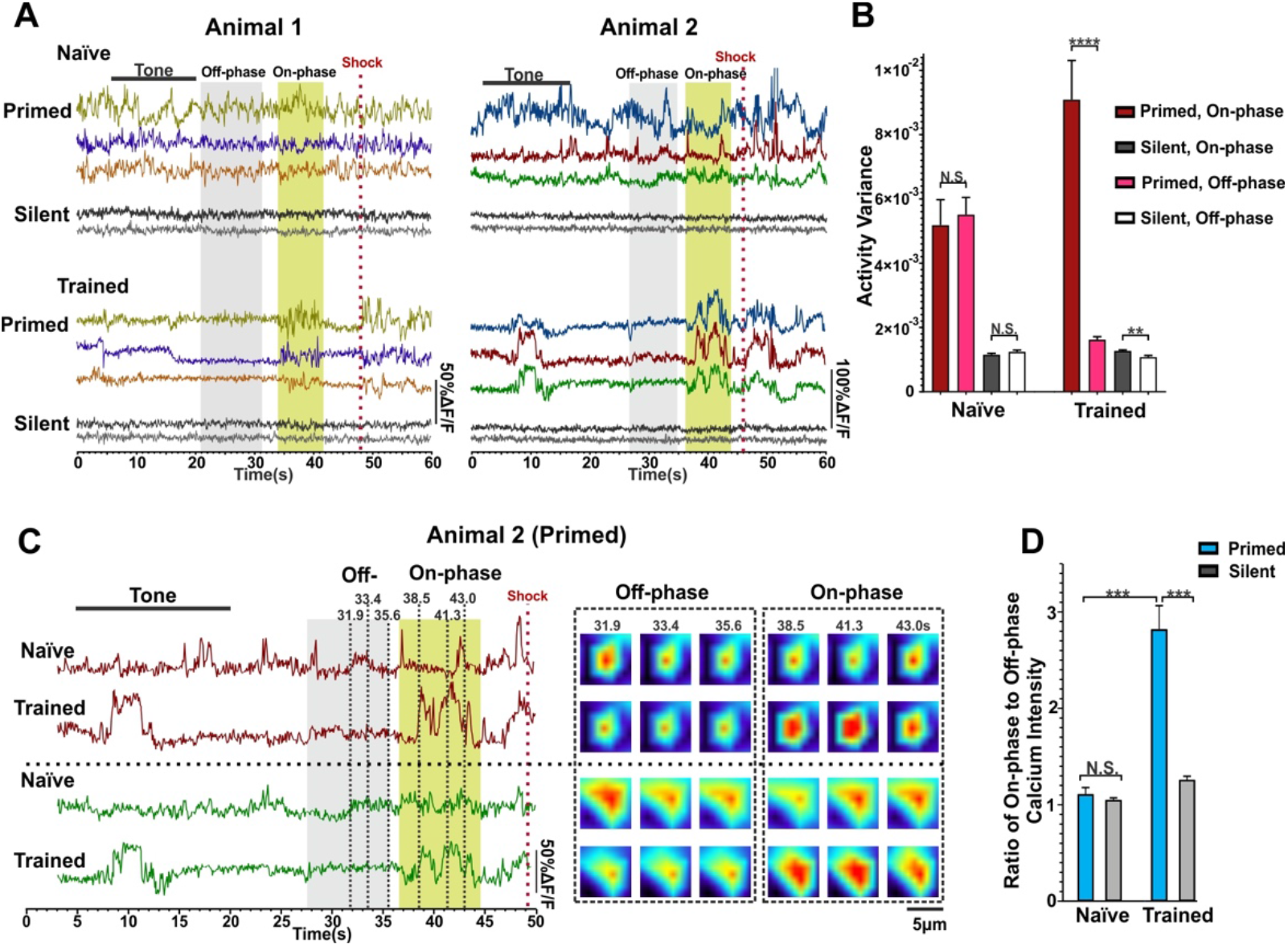
Training-induced synchronization condenses neuronal activity into a narrow time frame, promoting robust calcium entry into Primed Neurons. (A) Representative fluorescence traces of 3 Primed Neurons (color) and 2 Silent Neurons (grey) from two animals. Top, naïve condition; Bottom, after being trained. Low-activity “off-phase” and high activity “on-phase” periods are marked in gray and yellow shadow, respectively. (B) A bar graph presents the activity variance of on-phase and off-phase of Primed Neurons and of Silent Neurons under naïve condition and after being trained. Data were collected from 60 Primed Neurons out of 6 animals. Same number of 60 Silent Neurons were used as comparison. In naïve status, there were no differences in activity variance between on-phase and off-phase of Primed Neurons and of Silent Neurons. After being trained, Primed Neurons had increased on-phase activity variance and decreased off-phase activity variance. The activity variance of both on-phase and off-phase of Silent Neurons remained low. N.S., no significant difference; ***, p<0.001 by paired Student’s t-test. (C-D). The on-phase calcium entry of Primed Neurons drastically increased by CS-US paired training. (C) Left: fluorescence traces of 2 Primed Neurons from the animal 2 in naïve condition Vs. after being trained. Off-phase and on-phase periods are marked in gray and yellow shadow respectively. Right: representative images of 2 Primed Neurons in naïve condition versus after being trained. Dash black lines in the left traces shows the time points (during off-phase and on-phase period) when images were selected. (D) Bar graph shows the ratio of on-phase and off-phase calcium intensity. Calcium intensity areas were collected from 60 Primed Neurons and 60 Silent Neurons out of 6 animals. No all silent neurons were included in the variance calculation to avoid biased standard error of the mean. Naïvely, there were no differences in calcium intensity between on-phase and off-phase. After being trained, the ratio of on-phase and off-phase calcium intensity of Primed Neurons drastically increased, whereas that of Silent Neurons did not change much with training. N.S., no significant difference; ***, p<0.001 with paired Student’s t-test.

We compared on-phase and off-phase activity variance before training and after being trained. Prior to training, Primed Neurons’ calcium dynamics were irregular and had high activity variance in both on-phase and off-phase, suggesting that synaptic projects from the network onto these neurons were irregular, in line with a low synchrony index among Primed Neurons prior to training (Figure 5G). The on-phase to off-phase contrast became obvious after training (Figure 6A-bottom), due to increased on-phase activity and decreased off-phase activity (Figure 6A-B). Notably, after being trained, the off-phase activity variance of Primed Neurons was close to the “bottom” level of Silent Neurons’ activity variance (Figure 6B), while the on-phase activity variance increased by more than 50% (Figure 6B). These results suggest that CS-US paired training attuned Primed Neurons in the engram network to fire intensively during on-phase, but stayed temporarily silent during off-phase. In comparison, there was little change in calcium dynamics of Silent Neurons. Both their on-phase and off-phase signal variance (Figure 6A-B) and synchrony index and SNR (Figure 5G-H) remained low and did not change much with training, confirming that Silent Neurons somehow temporarily disengaged from the engram network.

How could activity synchronization affect trace memory formation? One possible impact is to condense the activity of the Primed Neuron population into a narrow time frame and intensify electrical stimulation onto postsynaptic neurons, which also include Primed Neurons themselves via recurrent synapses. This would strongly increase activity-associated calcium entry. Indeed, we found that prior to training, there were no differences in calcium dynamics to distinguish off-phase with on-phase activity (Figure 6C). The calcium dynamics of Primed Neurons were randomly distributed, and overall their intensity was not robust (Figure 6C). The ratio of on-phase to off-phase calcium intensity of both Primed Neurons and Silent Neurons was close to 1 (Figure 6C&D). Training-induced synchronization drastically increased the calcium intensity of on-phase of Primed Neurons, and decreased that of off-phase (Figure 6C&D). The ratio of on-phase to off-phase calcium intensity of Primed Neurons increased by ∼3 folds after being trained, while that of Silent Neurons did not alter much (Figure 6D). Note that after forming activity synchronization, on-phase synchronized activity could be induced by tone or spontaneously occur during the trace period (Figure 6A). The tone could sometimes (Figure 6C), but not always (Figure 6A), function as an inducer to trigger synchronized on-phase activity. Together, these data suggest that training-induced synchronization facilitates robust activity-associated calcium entry of Primed Neurons, which may subsequently send a strong signal to the nuclei to trigger gene expression to sustain long term potential and memory consolidation^39,40^.

## Discussion

This study has presented multiple exciting findings related to trace fear memory formation. First, the overall activity levels of principal CA1 neurons showed a right-skewed lognormal-like distribution. A small portion of Primed Neurons could maintain randomly active status for 5 hours in a day in multiple contexts, and short periods of sleep did not interrupt the activity hierarchy of CA1 neurons. The majority of CA1 neurons stayed silent during training and recall. Second, the highly active “Primed Neurons” were preferably selected to encode trace fear memory. Third, the activity dynamics of Primed Neurons were incrementally modified from a randomly active pattern to a well-tuned activity synchronization by repetitive CS-US paired trainings. Importantly, the emergence of synchronization correlated with the appearance of freezing behaviors in mice. Fourth, the activity dynamics of Primed Neurons were attuned by CS from chaotic activity to form partial synchronization, which also coincided with animal freezing behaviors. Additionally, training-induced synchronization intensified calcium entry into Primed Neurons.

Commonly used *in vivo* imaging techniques to visualize neuronal activity in the mouse brain are limited by several factors, including the need of a head-mounted microscope, severe tissue damage in the brain, imaging instability, and physical restriction on animals^41–43^. Use of a fiber optic confocal imaging technique confers several advantages. (1) The endoscopic imaging system does not restrict animal’s movement, allowing for imaging in more physiological conditions (Figure 1). (2) It is a minimally invasive surgery and imaging procedure, and animals can quickly recover from cannula implant surgery. (3) The *in vivo* imaging can be stably conducted, and we were able to record the activity of the same population of neurons up to 7 hours (Figures 1). We could successfully visualize the real-time dynamics of the same population of CA1 principal neurons that participate in learning and subsequent memory recall hours later. (4) We could simultaneously monitor the activity of a large population of neurons, typically 50-200 neurons. This allowed us to categorize neurons into different groups according to their activity features (Figure 3 and Table 1). Nevertheless, the fiber-optic confocal imaging system does have limitations. It has a limited spatial (3 µm) and temporal resolution (11.7 - 40 Hz). It cannot detect precise temporal information pertaining to neuronal activity during learning or resolve single action potential-caused calcium spikes. Moreover, imaging stability can be maintained in the same day, but not in the following days. In addition, imaging signals of this system are quite noisy and an effective method to reduce imaging noise is yet to be developed.

This *in vivo* imaging study in freely behaving mice has provided strong evidence supporting the concept that to participate in memory formation, hippocampal principal CA1 neurons need to be sensitized or primed. Primed Neurons demonstrated manifold advantages over Silent Neurons to be preferably recruited to encode trace fear memory. To encode memory information, memory-eligible neurons must be plastic to modify their activity pattern, and they can also easily change their activity pattern in response to memory cues to enable memory expression. Primed Neurons were sensitive to sensory stimuli and became more responsive to tone and foot shock after being trained (Figure 2). The activity of Primed Neurons was also highly plastic compared to Silent Neurons, allowing Primed Neurons to readily modify activity patterns in response to CS-US paired training or to CS in recall. In contrast, most CA1 neurons stayed silent in multiple contexts (Figure 3C). Little activities were evoked by tone or foot shock in learning and no event was detectable in Silent Neurons after repetitive foot shock stimulation. Hence, the contributions of Silent Neurons to memory formation may be very weak or. Our findings are consistent with recent reports that neurons in the amygdala^8^ and hippocampus^9^ with high excitability are preferably recruited to encode contextual fear memory^1,44–46^. Together, these findings strongly support a critical role of Primed Neurons in trace memory formation. Furthermore, this study further suggests that formation of declarative memory is governed by the availability of Primed Neurons prior to the training and by successful induction of synchronized activity among Primed Neurons. Likewise, retrieving a memory also depends on the availability of Primed Neurons prior to recall and whether synchronization among the same population of Primed Neurons can be effectively induced by memory cues.

Memory information is encoded and stored in the engram cell network^1,2,5^. Synchronized activity patterns in the neural network in neuronal culture models^47,48^ and theoretic computational models^49,50^ have been proposed as a reservoir of information^51,52^. However, direct *in vivo* evidence linking activity synchronization of hippocampal neurons with declarative memory formation was lacking. This study presents strong *in vivo* evidence, supporting a working model that trace fear conditioning facilitate forming a new neural circuit in the hippocampus, which wires up the CS-responsive neurons with the US-responsive neurons (Figure 7). Before training, Primed Neurons exhibit asynchronized activity, suggesting that there is no strong synaptic connection among these Primed Neurons. Increased activity synchronization among Primed Neurons reflects increased neural connectivity, based on the Hebbian Learning Rule that “cells that fire together wire together” ^4^. The newly formed synchronization among Primed Neurons suggests that they have built strong synaptic connections, likely via recurrent synapses among CA1 neurons^38^ or among the CA3 neurons^53,54^ that can further project to the CA1 neurons. Hence Primed Neurons can fire action potentials in high coherence. The new connecting circuit is incrementally constructed, and their synaptic connection becomes increasingly stronger with repetitive training (Figure 7). After training, although synchronization can persist for a short while, there is eventually a return to chaotic dynamics, which may allow for encoding new information. Prior to the subsequent recall, Primed Neurons are randomly active, but they can readily modify their activity pattern from a random to a synchronized mode in response to a cue. If a strong synaptic circuit that electrically connects Primed Neurons has been constructed, activation of a few neurons or even a single neuron (e.g., a tone-responsive neuron) in the network could markedly impact the dynamics of the network^55,56^. It induces partial synchronization, which could be sufficient to electrically connect the tone-responsive neurons with the foot shock-responsive neurons, consequently causing freezing responses (Figure 7).

**Figure 7.**
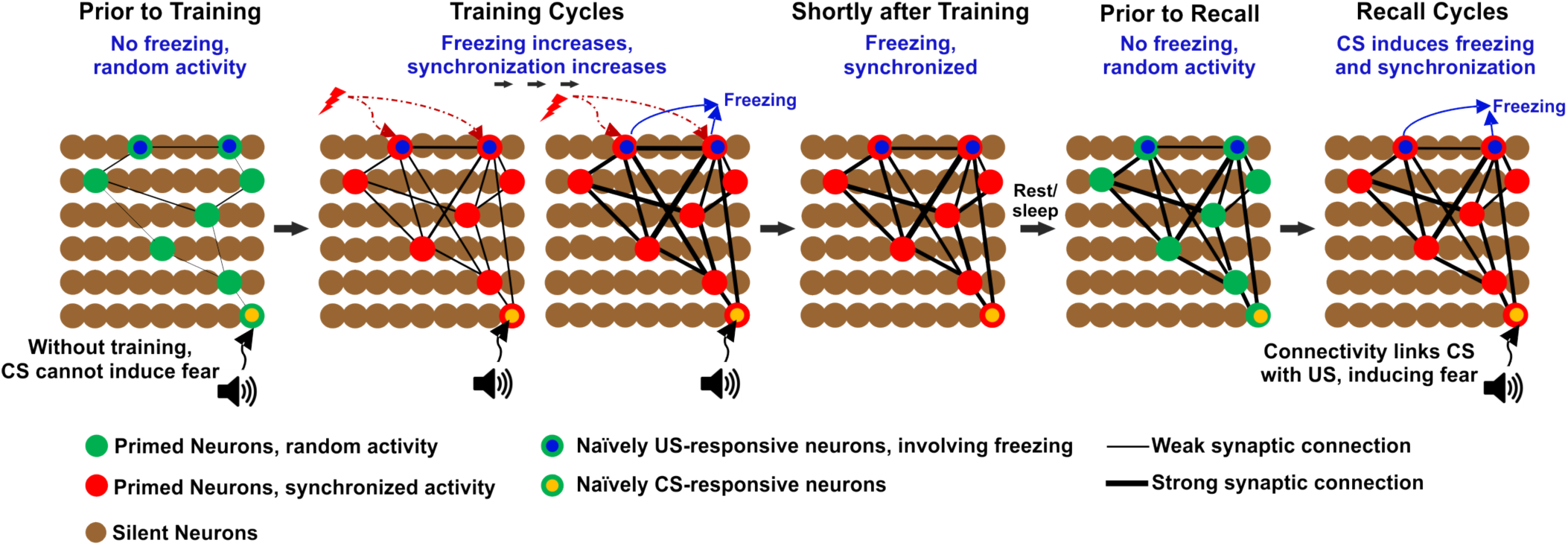
A tentative model for trace memory formation: Primed Neurons forge a new circuit to bridge CS with US through connecting with one another. Green: Primed Neurons with random activity. Red: Primed Neurons with synchronized activity during training and recall testing. Brown: Silent Neurons. Blue dot in green cells: naïvely foot shock-responsive neurons, whose activation is naïvely associated with freezing responses (US-responsive neurons). Yellow dot in green cells: naïvely tone-responsive neurons, whose activation does not cause fear responses (CS-responsive neurons). Multiple cycles of CS-US paired training induced activity synchronization among Primed Neurons, suggesting that the connectivity among Primed Neurons increases. After Primed Neurons have connected with each other and forged a new circuit, stimulation of a few CS-responding neurons would markedly impact the network dynamics and induce synchronized activity in the network. This could consequently activate US-responsive neurons, leading to freezing responses. Note that the newly formed synaptic connection could be directly occurring in the CA1 region via recurrent synapses^38^, or indirectly involving the hippocampal CA3 region^53,54^ and then project to CA1. Intermediate Neurons exhibit mild activity and may follow Primed Neurons to participate in memory formation. Nevertheless, they are not included in this model for simplicity.

It is worth mentioning that the first impact of activity synchronization on memory formation is that synchronized neuronal activity in the network, in which all Primed Neurons fire in a narrow time frame in a bursting pattern, promotes marked elevation of intracellular calcium during learning (Figure 6). Consequently, it could trigger gene expression to potentiate and sustain high synaptic efficacy for memory consolidation^39,40^. Second, activity synchronization may also affect memory formation when Primed Neurons intertwined with recurrent synapses fire with a bursting pattern, since almost at the same time they also receive coherent synaptic input from other Primed Neurons, because other neurons in the engram network also fire. This would lead to high-frequency electric stimulation on individual postsynaptic neurons. Third, because of the activity synchronization and intertwined synaptic connections, the action potential events of Primed Neurons and their synaptic activities in the dendrites can occur within a narrow time frame, which may enable spike-timing-dependent plasticity^57–59^ to enhance the efficacy of synaptic transmission among Primed Neurons.

Our findings drastically modify a prevailing model of memory formation, in which a sparsely distributed population of engram cells in memory-related regions^60^ “activates” during learning to encode a memory and the same population of neurons “re-activates” to retrieve the memory^7^. Our data clearly show that memory-eligible Primed Neurons are already highly active prior to training, but their activities display random dynamics and have low synchronization prior to training. Primed Neurons change their activity pattern in response to CS-US paired trainings, develop activity synchronization and form a neural circuit. Primed Neurons also stay randomly active prior to the recall, and CS induces activity synchronization to retrieve a memory. In contrast, Silent Neurons are always silent throughout the training cycles, even when stimulated with multiple foot shocks. This suggests that it is difficult to directly activate the Silent Neurons to become active and to participate in memory formation, and neurons need to be first primed to become memory-eligible to participate in memory formation. Additionally, the synchronization of Primed Neurons also is correlated with animals’ freezing behaviors. Together, our findings suggest that the proposed “neuronal activation” for memory formation does not mean that neuronal activation arises from a completely silent state. More precisely, it is the activity dynamics of Primed Neurons that are subjected to modification from asynchrony to synchrony in learning to encode memory information. Moreover, memory retrieval is not simply the “re-activation” of engram cells from their silent state. Rather, it is the same population of memory-holding Primed Neurons, which are randomly active prior to the recall, that change their activity pattern to form activity synchronization to retrieve a memory.

In conclusion, trace fear memory is preferably encoded by Primed Neurons, whose dynamics are shifted from random activity to form activity synchronization during training. In recall testing, CS-induced activity synchronization among the same group of Primed Neurons helps retrieve the memory.

## Acknowledgements

We thank Drs. Zhengui Xia, Dan Storm, and Larry Zweifel for research and technical support. We are grateful to Dr. Rick Cote and the members of Chen Laboratory for their critical reading of the manuscript. This study was supported by National Institutes of Health Grants MH105746, AG054729, and GM113131-7006 as well as Cole Neuroscience Faculty Research Awards to X.C.

## Legends for Supplemental Files

**Supplemental Video 1**: A video of calcium imaging time-locked with mouse behavior at the first training (Cy1, naïve) of trace fear conditioning. The top left window shows a full endoscopic imaging video with 3 Primed Neurons (a, b, c) and 3 Silent Neurons (1, 2, 3) labeled *in situ*. The top-right window shows mouse behavior. The mouse has not associated tone with foot shock. The real-time calcium fluorescence traces (%ΔF/F) of 3 Primed Neurons (a, b, c) and 3 Silent Neurons (1, 2, 3) are shown in the bottom. All neurons had the same scale bars.

**Supplemental Video 2**: A video of calcium imaging and mouse behavior at the 7^th^ training cycle (Cy7, being trained) of trace fear conditioning. The top left window shows a full endoscopic video with the same 3 Primed Neurons and 3 Silent Neurons labeled *in situ*. The top right window shows animal behaviors. After 7 cycles of training, the mouse has associated tone with aversive foot shock stimuli. Traces of 3 Primed Neurons (a, b, c) and 3 Silent Neurons (1, 2, 3) are shown in the bottom. All neuronal traces had the same scale bars.

**Supplemental Video 3**: A video of imaging when the mouse was sleeping. The same 3 Primed Neurons (a, b, c) and 3 Silent Neurons (1, 2, 3) are labeled. During sleep, both Primed Neurons and Silent Neurons were not active. The three videos were acquired from the same animal, from which traces of the 3 Primed Neurons and 3 Silent Neurons were derived.

**Supplemental Figure 1:**
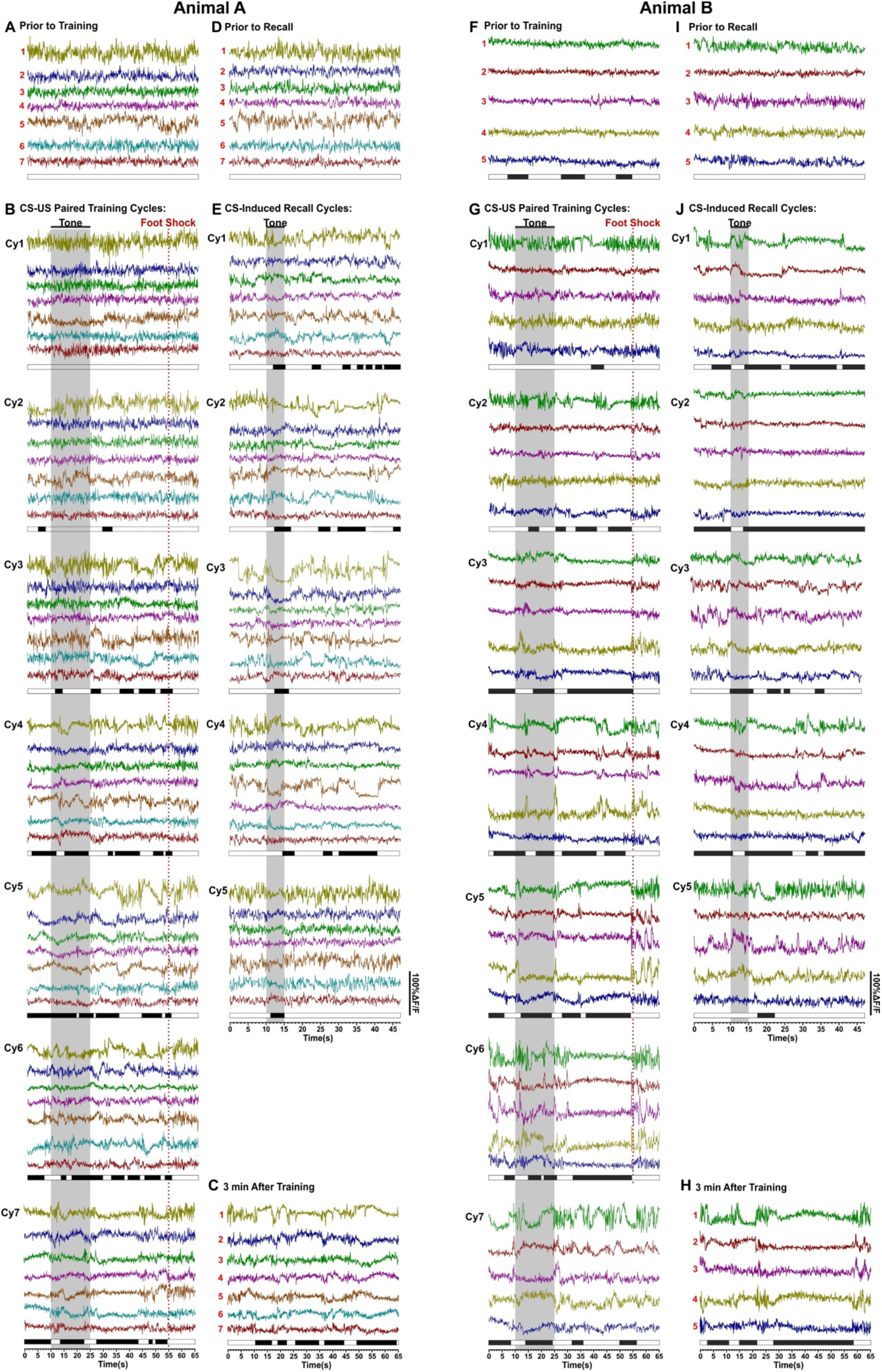
Primed Neurons modify their activity pattern to form activity synchronization during trace fear conditioning and recall. Traces of Primed Neurons from two animals are shown. (A-E) Traces of 7 Primed Neurons from one animal, which has been partly shown in Figure 5. White bars and black bars underneath the traces show animal moving and freezing, respectively. (A) Prior to training. (B) 7 training cycles. (C) 3 min after training. (D) Three minutes prior to the recall testing. (E) Recall testing cycles. (F-J) Representative fluorescence traces of 5 Primed Neurons acquired from another animal. Prior to training (F), during training (G), after training (H), Prior to recall (I), and during recall testing (J). Note that the Animal B was not very active or had spontaneous freezing prior to training (F), which made Primed Neurons less active. In Cy1, the animal became normally active. In Cy7, a foot shock stimulus failed to be efficiently delivered to the animal.

